# Temporal and Spatial Niche Partitioning of Bumblebee Species (Apidae: *Bombus*) on a Himalayan Mountain

**DOI:** 10.1101/2021.06.30.450469

**Authors:** Alan James Moss, Gerardo R Camilo, Zong-Xin Ren, Peter Bernhardt

## Abstract

**Background:** Bumblebees are essential pollinators in many ecosystems. To maintain such ecosystem functions, bumblebee diversity remains an important concern. We measured abundance and diversity of bumblebee species along an elevational gradient over three flowering seasons. We hypothesized that co-occurring bumblebees partition niches over time and space.

**Methods:** We sampled bumblebees at six different meadows along Yulong Snow Mountain (Yunnan, SW China), a region known for high bumblebee diversity, over summer months (2017 – 2019). We analyzed the standardized abundances of the workers from different bumblebee species over 11 weeklong periods. We also analyzed the standardized abundances of the workers of each species at each of the six different meadows.

**Results:** Clear patterns of temporal niche partitioning were most apparent for the two, dominant species, *B. friseanus* and *B. lepidus*. Spatial niche partitioning was evident for rarer species which tended to be most abundant at lower elevations.

**Conclusions:** Temporal and spatial niche partitioning are among two of the ways that bumblebee diversity is maintained on Yulong Snow Mountain. This has future implications if changing climate conditions disrupt partitioning leading to increased interspecific competition.

## Introduction

Bumblebees (*Bombus* Latreille) are important pollinators in numerous ecosystems and for some commercial crops (Corbet, Williams and Osborne 1992; Goulson 2010; Kevan 1991; Memmott, Waser and Price 2004). As the eusocial behavior of gynes and workers requires they collect enough pollen and nectar to feed growing colonies (Heinrich 1979), this leads to longer foraging bouts and additional opportunities for conspecific pollen transfer increasing cross-pollination events (Pleasants 1980). In addition, bumblebees are also uniquely adapted for surviving at higher elevations due to thermoregulatory traits including large hairy bodies that can efficiently retain heat produced by vibration of their large wing muscles (Ollerton 2021; Heinrich 1974). Consequently, bumblebees are among the most common pollen vectors on mountain slopes and peaks with many species of temperate and tropical montane plants (Bernhardt and Montalvo 1977) dependent on them for pollination services (Macior and Tang 1997).

Although bumblebee species are distributed on most continents, their center of diversity is within the Eastern Himalayas where half of the world’s estimated 250+ species live (Huang and An 2018; Williams 1998; Williams et al. 2009). While much is known about regional bumblebee diversity (i.e. gamma diversity *sensu* Whittaker 1960) much less is known about how local bumblebee diversity (alpha diversity) is organized and how species turnover (beta diversity) affects bumblebee distributions. In some regions, several bumblebee species all forage in the same meadow (Goulson 2010). In particular, study sites located on Yulong Snow Mountain in Lijiang County (Yunnan, China) currently record 21 different bumblebee species (Williams et al. 2018, unpublished data). Adding to potential confusion, most bumblebee species are morphologically similar and often consume the same floral resources (Macior 1974; Pleasants 1980; Liang et al. 2018). For decades, the old question remains how do different bumblebee species coexist without significant competition (Inouye 1978)?

Extreme abiotic factors typify the Himalayas as air temperature and air pressure decrease with increasing elevation while wind velocity and solar radiation increase (Barry 2008). However, while increasing temperature is broadly associated with increasing species richness (Hawkins et al. 2003), patterns of species richness on mountains do not always follow this trend. Species richness on mountains tends to be either highest at the lowest elevations or it peaks along mid-montane elevations (McCain and Grytnes 2010). While species richness patterns of montane insects have been controversial (McCoy 1990) it is unlikely that abiotic factors alone determine the distributions of cold-hardy bumblebees. Bumblebees in the Himalayan region are often found at far higher elevations (Williams et al. 2010) compared to those reported in North America (Macior 1974). Some species are capable of flight in conditions simulating 9000 m above sea level (Dillon and Dudley 2014), higher than Mount Everest at 8,840 m (Stegman 1982). Consequently, with numerous adaptations to alpine conditions, bumblebee diversity is more likely constrained by additional factors.

The argument for biotic factors driving bumblebee distributions is based on the competitive exclusion principle (Hardin 1960) in which two species occupying the same ecological niche end eventually in the local extinction of one species due to exclusionary competition by the second. Numerous bumblebee species occurring in the same meadow on the same mountain would appear to violate this ecological principle. The most contested resource for eusocial bumblebees must be floral resources with nectar as a food for adults while a combination of pollen and nectar is fed to larvae. These insects have been shown to change their floral preferences based on the presence of sympatric congeners. Therefore, floral resources are a source of competition among different bumblebee species (Inouye 1978) just as montane plants compete for the limited resource of bumblebee pollinators (Pleasants 1980). Without changes limiting direct interspecific competition the Himalayan region of Yunnan could not be a bumblebee “hotspot.” Instead, there would be fewer species with each restricted precisely to elevation zones in which they were best adapted.

Instead, variation in life-histories make it possible for closely related but sympatric species to decrease competition and avoid extinction although their ecological niches overlap. For example, respective phenology may diverge significantly leading to differing peak periods in which two or more species are most abundant and are exploiting similar resources. This is known as temporal niche partitioning (Winder 2009). Staggering emergence times for sympatric bumblebee species should be very important as foraging must correspond to peaks of limited floral resources decreasing under interspecific competition (Goodwin 1995). This can also have repercussions for rates of plant fertilization and subsequent seed-set (Kudo and Cooper 2019; Memmott et al. 2007; Pleasants 1980). If bumblebees emerge in succession during the same season within the same elevational zone we should expect that their combined total abundance would remain stable linked to the relative abundance of flowering species with staggered blooming in succession.

Spatial niche partitioning is a second alternative. Interspecific competition is avoided but at a much finer scale. The classic example is of birds in the same lineage foraging in different layers within the same coniferous forest (MacArthur 1958). While spatial niche partitioning is documented in different species assemblages (Albrecht and Gotelli 2001; Buckley and Roughgarden 2005; Winder 2009; Zhong et al. 2016) the line could be blurred between abiotic and biotic influences affecting bumblebee distributions. To avoid direct interspecific competition, bumblebee species may have different thermoregulatory adaptations that permit specialization along narrowing elevation gradients.

Based on these potential components, the goal of this study was to determine if bumblebee species were spatially and/or temporally niche partitioned leading to lower interspecific competition within a hotspot in a section of the Chinese Himalayas. Our study asks two overlapping questions*. First, were there significant temporal variations in the foraging periods of workers of different bumblebee species and their relative abundance throughout the summer? Second, were there significant spatial niche partitionings of bumblebee species along an elevational gradient?* In the absence of either mitigating factor, significant differences in relative abundances between bumblebee species temporally and spatially would support a conclusion that interspecific competition leads to species richness losses.

## Materials & Methods

### Sample Sites Overview

Yulong Snow Mountain is part of the UNESCO World Heritage Site known as The Three Rivers Region of Yunnan, China. It is a biodiversity hotspot with the Yulong Snow Mountain supporting 2815 angiosperm species according to a checklist compiled by Liu et al. (2015). Portions of Yulong Mountain are under the management of the Lijiang Alpine Botanic Garden and the Lijiang Forest Ecosystem Research Station which is part of the Kunming Institute of Botany, Chinese Academy of Sciences (Kunming, Yunnan). The mountain is part of an expanding tourist industry and is a source of wild resources (edible plants and fungi, medicinal herbs, decorative timbers etc.) for many of the local peoples. Its meadows are traditional long-term grazing sites for domesticated yaks, goats, cattle, mules, and horses. These ecological services are culturally important for several indigenous minorities including the Naxi (Zhang et al. 2015), the largest minority group in the region.

Specifically, “Yulong Snow Mountain” is the name given by local people although it is a mountain chain of several peaks. The base of the specific mountainous region where we sampled was at approximately 2700 m above sea level [27°, 100.2°] (Fig. 1 A) while the highest peak we used for field collections was at 3900 m above sea level [27.02°, 100.18°] (Fig. 1 D).

**Fig. 1.**
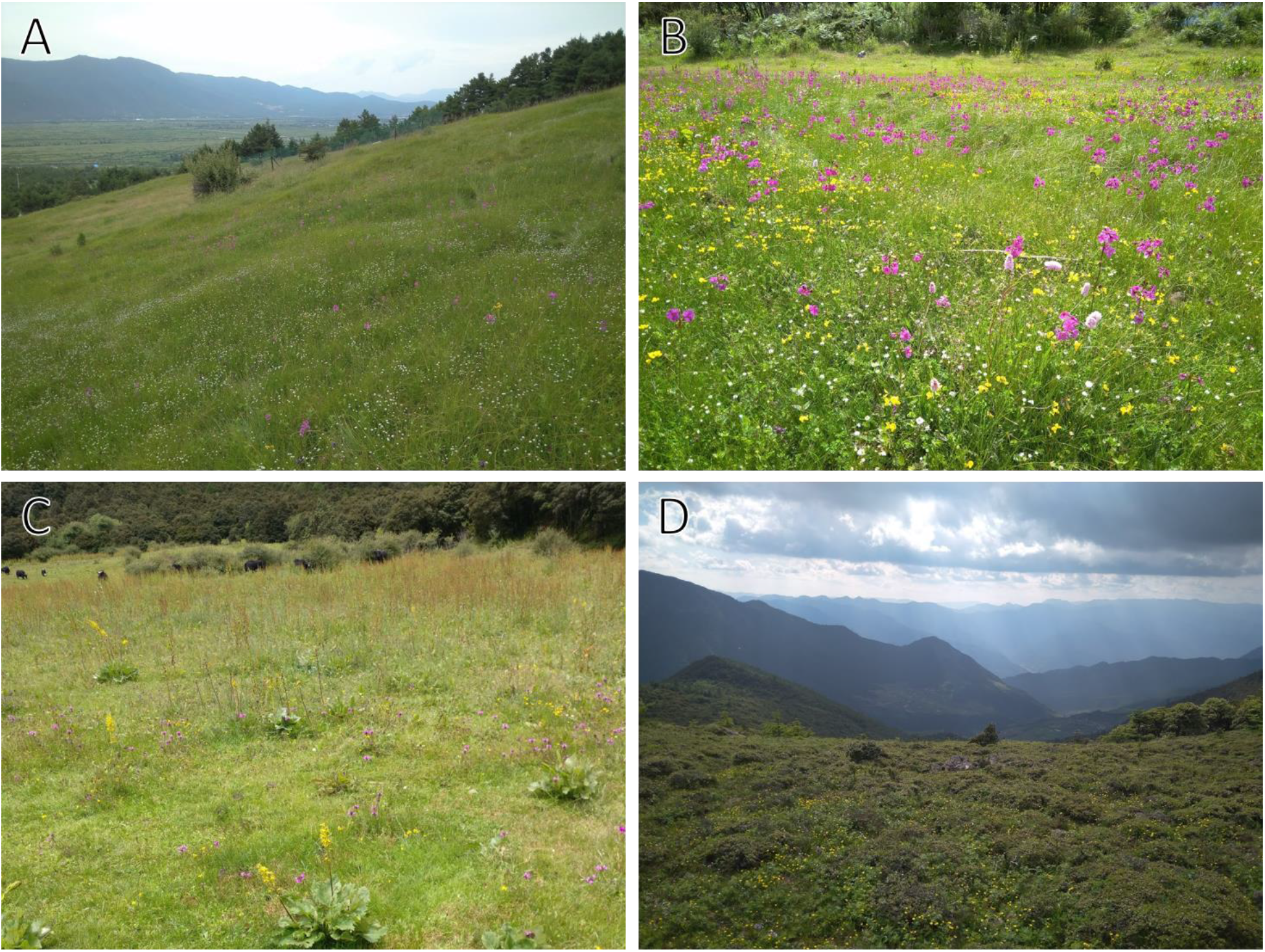
(a) Sample site, 2700 m, (b) sample site, 3200 m, (c) sample site, 3400 m, (d) sample site at summit, 3900 m

Most of Yulong Snow Mountain is covered in conifer forests with an understory of *Quercus* species (Luo et al. 2019). We restricted our study sites to wet meadows between or below forests and dry meadows located above forests (Fig. 1). Wet meadows are natural drainage sites of varying sizes found largely between peaks. The soil in wet meadows is too hydrated to support succession by trees. Dry, upland meadows are confined to peaks where cold temperatures, high winds and rocky extrusions also prevent tree succession. Despite the harsh abiotic conditions, the meadows are home to many bumblebee-pollinated species in the same genus (Liang et al. 2018) often occurring sympatrically as complicated species flocks (*sensu* Lecointre et al. 2013).

As Yulong Snow Mountain has numerous meadows, we selected the largest at median sizes of 33,950 meters squared (Table 1). We made this selection because large meadows tended to have less extreme gradients and fewer confounding factors such as a high edge-to-total-area ratio. Smaller meadows in contrast tended to be very narrow (in between conifer forests) with steep inclines. The dry meadow at the summit was the largest continuous meadow so we sampled only a portion of its total area. Specifically, we sampled this meadow at its highest elevation, roughly equivalent to the median size of the other meadows at lower elevations. The choice to sample only a portion of this summit meadow was made in part due to limitations imposed by steep gradients making sampling dangerous and because we wanted to sample a portion of the summit showing elevation homogeneity (see above). Bumblebee foraging distances vary with species. Some can forage several thousand meters from their nesting sites although the mean foraging distance is usually under 500 meters (Osborne et al. 1999; Redhead et al. 2016). A study by Geib, Strange and Galen (2019) of four alpine bumblebee species and their respective foraging distances most overlaps with the conditions of our study region. The authors found that the upper foraging range for their bumblebee species was under 300 meters. The closest of our sites were 930 meters apart, therefore it is unlikely that significant numbers of bumblebees foraged at more than one of our study sites on the same day.

**Table 1.**
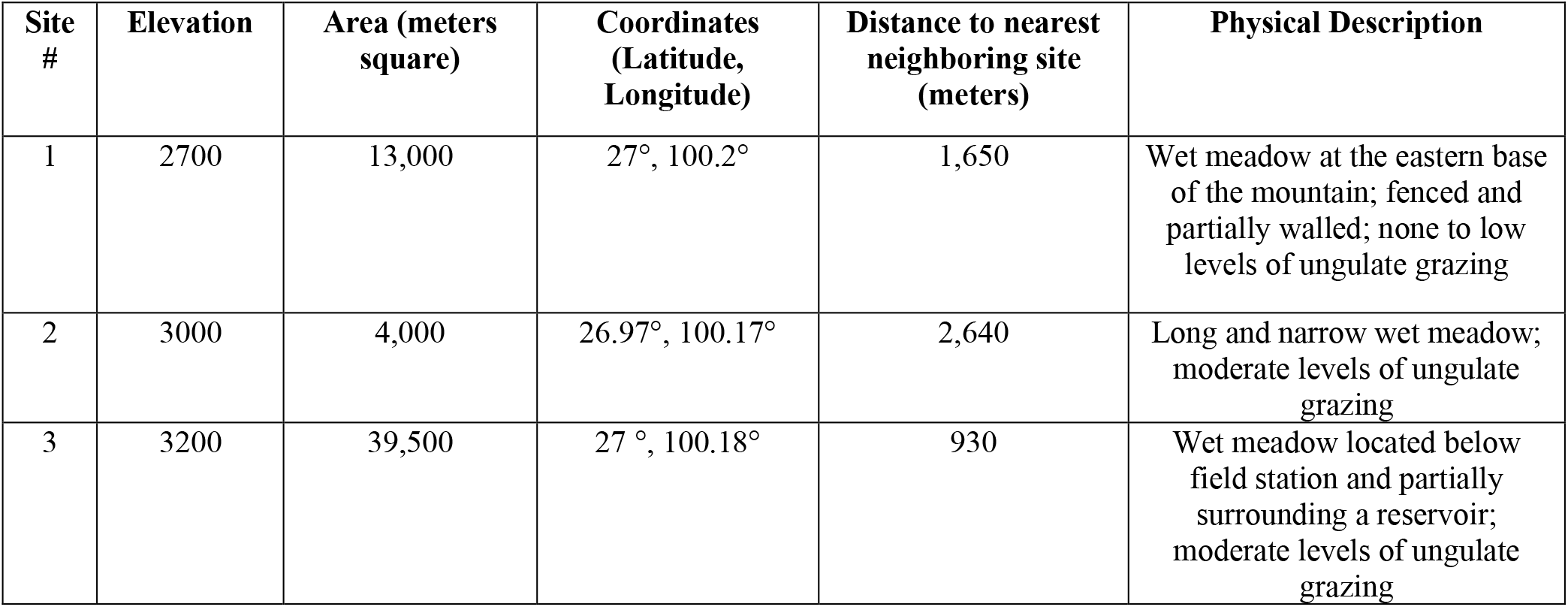
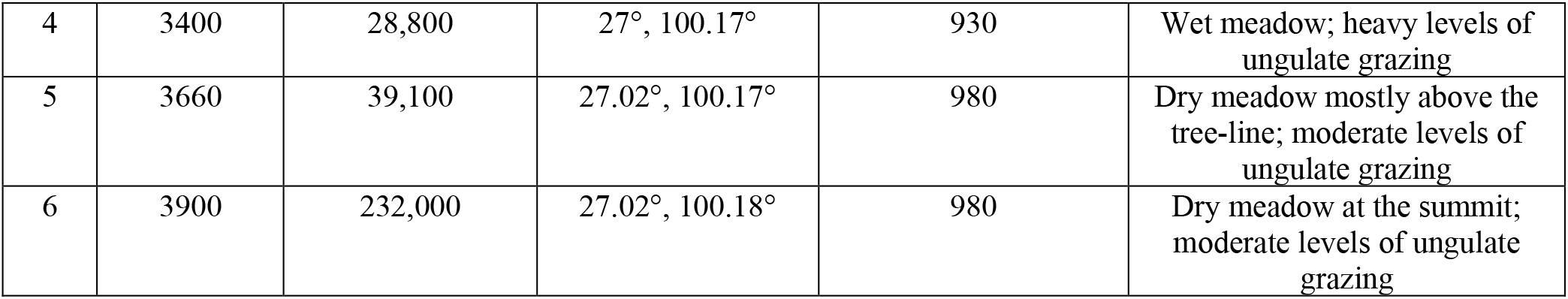
Yulong Snow Mountain sample site descriptions

| Site # | Elevation | Area (meters square) | Coordinates (Latitude, Longitude) | Distance to nearest neighboring site (meters) | Physical Description |
| --- | --- | --- | --- | --- | --- |
| 1 | 2700 | 13,000 | 27°, 100.2° | 1,650 | Wet meadow at the eastern base of the mountain; fenced and partially walled; none to low levels of ungulate grazing |
| 2 | 3000 | 4,000 | 26.97°, 100.17° | 2,640 | Long and narrow wet meadow; moderate levels of ungulate grazing |
| 3 | 3200 | 39,500 | 27 °, 100.18° | 930 | Wet meadow located below field station and partially surrounding a reservoir; moderate levels of ungulate grazing |
| 4 | 3400 | 28,800 | 27°, 100.17° | 930 | Wet meadow; heavy levels of ungulate grazing |
| 5 | 3660 | 39,100 | 27.02°, 100.17° | 980 | Dry meadow mostly above the tree-line; moderate levels of ungulate grazing |
| 6 | 3900 | 232,000 | 27.02°, 100.18° | 980 | Dry meadow at the summit; moderate levels of ungulate grazing |

### Sampling Regime

Yulong Snow Mountain undergoes a yearly monsoon season centered in the months of June-August during which time rain falls most days (Thomas 1993). This monsoon period also overlaps with the peak flowering season when most bumblebee species are active. Rain was a constant complication for sampling as it made standardizing collection efforts difficult. Weather in the Himalayas is often unpredictable changing quickly and making it perilous to sample. Therefore, we often had to cancel or interrupt sampling at field sites during periods of heavy or severe rainfall. To standardize collection efforts, bumblebees were captured for approximately 1 hour per site. Due to frequent interruptions by unpredictable rains, we often had to remain at each site for 4 or more hours, collecting bumblebees for short intervals between bouts of rain and fog until the total actual collection time pooled to 1 hour. Bumblebees were collected exclusively during daylight hours avoiding the time around solar noon (12 - 2 PM) when bumblebee flight activity declines due to increased solar radiation (Xu et al. 2021, unpublished data).

Collections were made from June 24^th^ to July 26^th^ in 2017, from August 7^th^ to September 3^rd^ in 2018, and from July 7^th^ to August 8^th^ in 2019. Research sites at 2700m, 3200m, and 3400m were sampled all three years. The 3000m site was sampled in 2018 and not used again because of low insect activity. The 3660m and 3900m sites were added in 2019. These two highest sites where largely inaccessible the two previous years of 2017 and 2018, due to heavy rains, but were added in 2019 because drought made the dry meadows near the peak accessible to sampling.

Each site was sampled weekly via aerial netting. Bumblebees were netted only when they were observed foraging actively for nectar and/or pollen. Collected specimens were put into individual tubes to avoid cross contamination of pollen loads for future analyses. Several tubes of different sizes were tried but the most effective was the 5mL screw cap tube (VWR catalog number: 470225-026) because it easily held a single bumblebee but did not take up excess space in field bags or in the freezer. Specimens were euthanized at the KIB field station (3200m).

### Processing of Bumblebee Specimens

Identification to species followed Williams (2018, unpublished). Bumblebees were pinned vertically, labeled, and are stored as vouchers in the Pollination Laboratory of the Kunming Institute of Botany, Chinese Academy of Sciences.

### Data Analysis

Only female workers were included in analyses to avoid potential confounding effects from different castes. The periods of sampling over three summers when pooled led to a greater sampling effort in the middle of the summer and fewer samplings toward the beginning and ends of summer. To account for the unevenness of sampling effort, the combined summer collection period was split into 11 time periods. Each period was 7 days long, from June 24^th^ to September 8^th^. The total abundance of each species collected at each site was then divided by the number of times that site had been sampled during that week period over the three summers (see supplemental [datasheet]) with standardized abundances. All statistical analyses were done using R (R Core Team 2020), a statistical computing language in the integrated development environment of RStudio (RStudio Team 2020) and using the data visualization package ‘ggplot2’ (Wickham 2016).

### Temporal Analysis

We compared and contrasted the flight/foraging period of each bumblebee species via visual representation. We created a bar plot with an overlaid smoothed conditional mean line using the “loess” method which showed the standardized abundance of each species over 11 week-long periods. To better compare the two most dominant species, with dominance determined by relative abundance, we created a second bar plot with an overlaid, smoothed, and conditional mean line. We then used a boxplot graph to show the distribution of each species over time. Finally, we calculated the weighted mean week of occurrence by taking the mean of all weeks, numbered 1 through 11, when each species was collected weighted by its standardized abundance for each week. Weighted mean was found by Moussus, Julliard and Jiguet (2010) to be a more accurate estimate of phenophase than most other measures such as first recorded occurrence.

### Elevational Distribution Analysis

We analyzed spatial patterns, to understand relative abundances of each species along an elevational gradient. First, we visualized the distributions of each species using a bar plot graph with an overlaid, smoothed, and conditional mean. We then made a boxplot graph to further visualize the elevational distribution of different bumblebee species, as above.

## Results

Over three summers of sampling on Yulong Mountain, 14 different species of bumblebees were identified. Of these, five species [*B. funerarius* Smith, 1852; *B. keriensis* Morawitz, 1887; *B. religiosus* Frison, 1935; *B. securus* (Frison, 1935); *B. waltoni* Cockerell, 1910] were excluded from further analyses as we caught less than five specimens of each taxon. This left a total of 531 worker specimens that were used in the analysis. Bumblebee worker abundances, elevational ranges, weighted mean week of occurrence, and active flight periods are summarized in Table 2.

**Table 2.** Correlation of *Bombus* species collection periods, their elevation ranges, phenology, and flight periods

| *Bombus* sp. | Number of Workers Collected | Elevational Range (meters) | Weighted Mean Week of Occurrence | Worker Flight Period |
| --- | --- | --- | --- | --- |
| *avanus*<br>(Skorikov, 1938) | 5 | 3200 – 3400 | 6.07 | 25 July – 7 Aug. |
| *festivus*<br>Smith 1861 | 8 | 2700 – 3200 | 6.2 | 8 July – 14 Aug. |
| *friseanus*<br>Skorikov, 1933 | 335 | 2700 – 3900 | 6.61 | 26 June – 3 Sept. |
| *grahami*<br>(Frison, 1933) | 7 | 2700 – 3200 | 6.51 | 2 July – 2 Sept. |
| *impetuosus*<br>Smith, 1871 | 7 | 2700 | 2.61 | 2 July – 14 July |
| *infrequens*<br>(Tkalcu, 1989) | 17 | 2700 – 3660 | 5.96 | 8 July – 20 Aug. |
| *lepidus*<br>Skorikov, 1912 | 150 | 2700 – 3900 | 3.28 | 24 June – 18 Aug. |
| *longipennis*<br>Friese, 1918 | 5 | 3200 – 3900 | 2.88 | 2 July – 18 July |
| *remotus*<br>(Tkalcu, 1968) | 9 | 2700 – 3400 | 1.55 | 5 July – 26 July |

### Temporal Patterns

The standardized abundance of the remaining nine species over an 11-week period is shown in Figure 2. Two species, *B. friseanus* and *B. lepidus*, were the most abundant (Table 2). *Bombus friseanus* and *B. lepidus* showed flight/foraging activity for most of the 11-week period of sampling. Remaining species showed activity for shorter periods. A total of 323 workers were collected of *B. friseanus* and 150 for *B. lepidus.* The next most abundant species was *B. infrequens* (n = 17 specimens). Clearly *B. friseanus* and *B. lepidus* were the dominant species and were potential competitors of each other. However, *B. lepidus* was more abundant in early summer while *B. friseanus* was more abundant later in the season (Fig. 3). Figure 4 shows the distribution when each species was active.

**Fig. 2.**
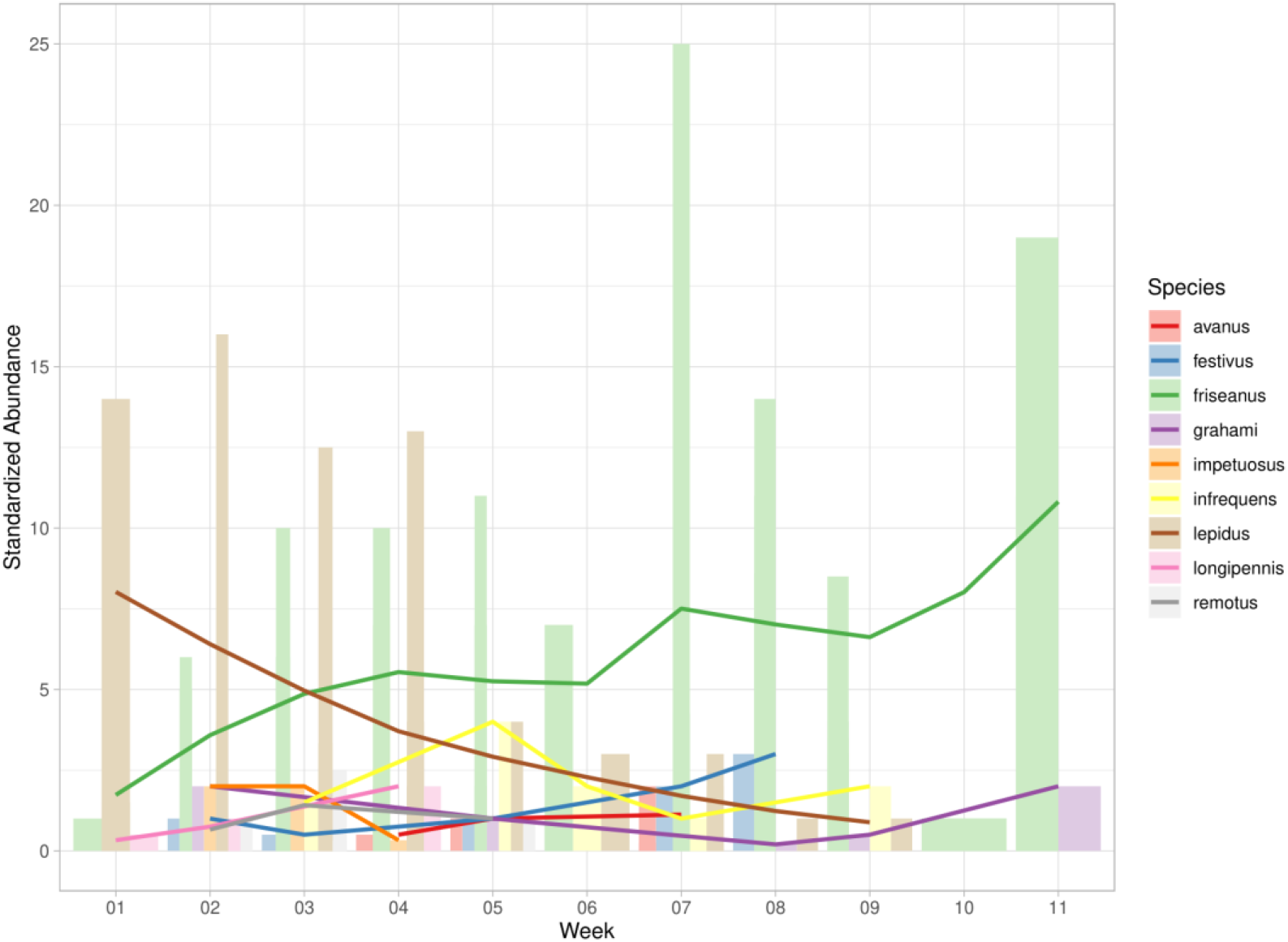
Comparative abundances, standardized by collection effort, for bumblebee (*Bombus*) workers over 11 week-long periods. Lines indicate the smoothed conditional mean of abundance for each species

**Fig. 3.**
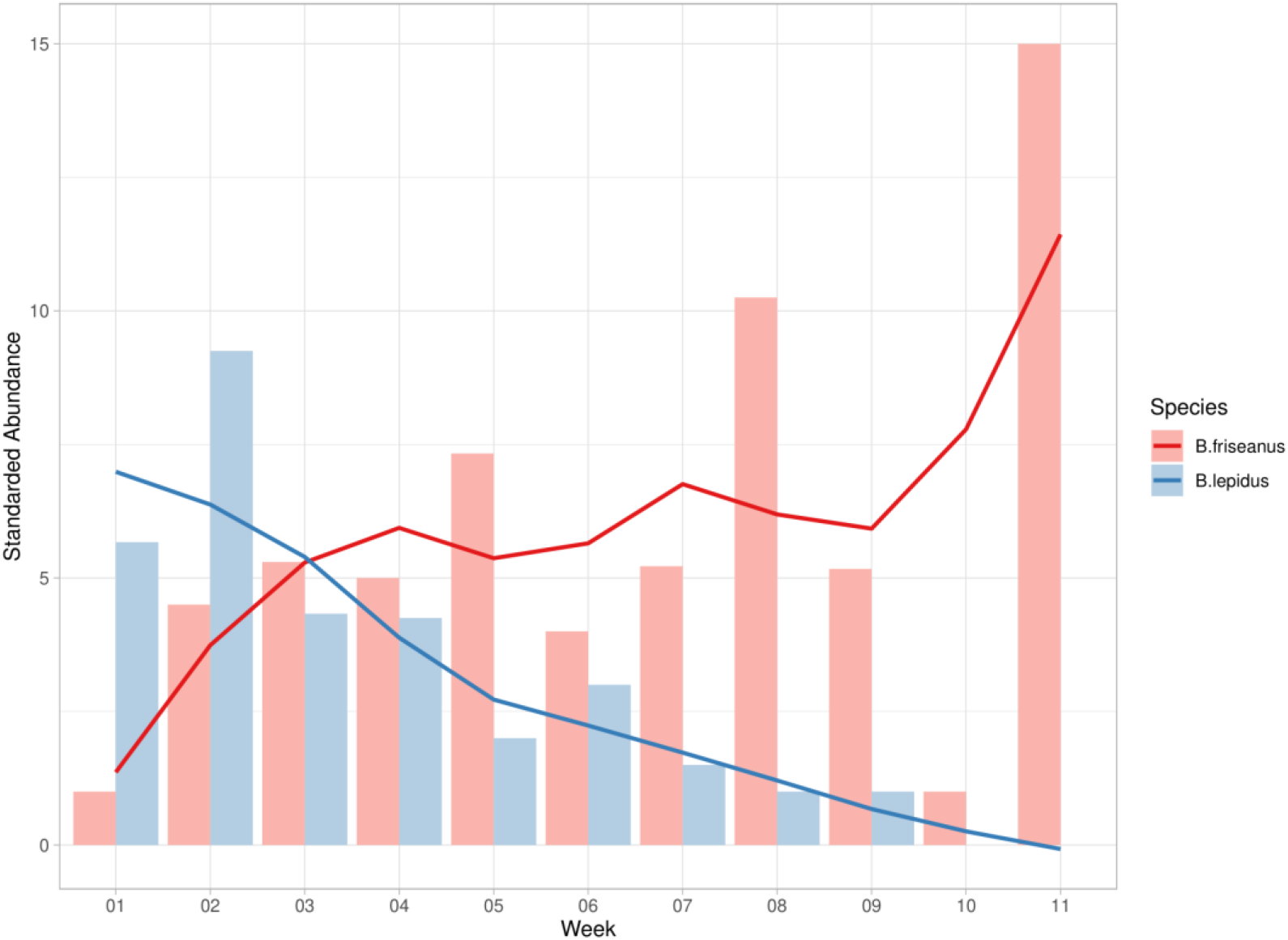
Standardized abundances of workers with smoothed conditional mean lines for *Bombus friseanus* and *B. lepidus* over 11 week-long periods

**Fig. 4.**
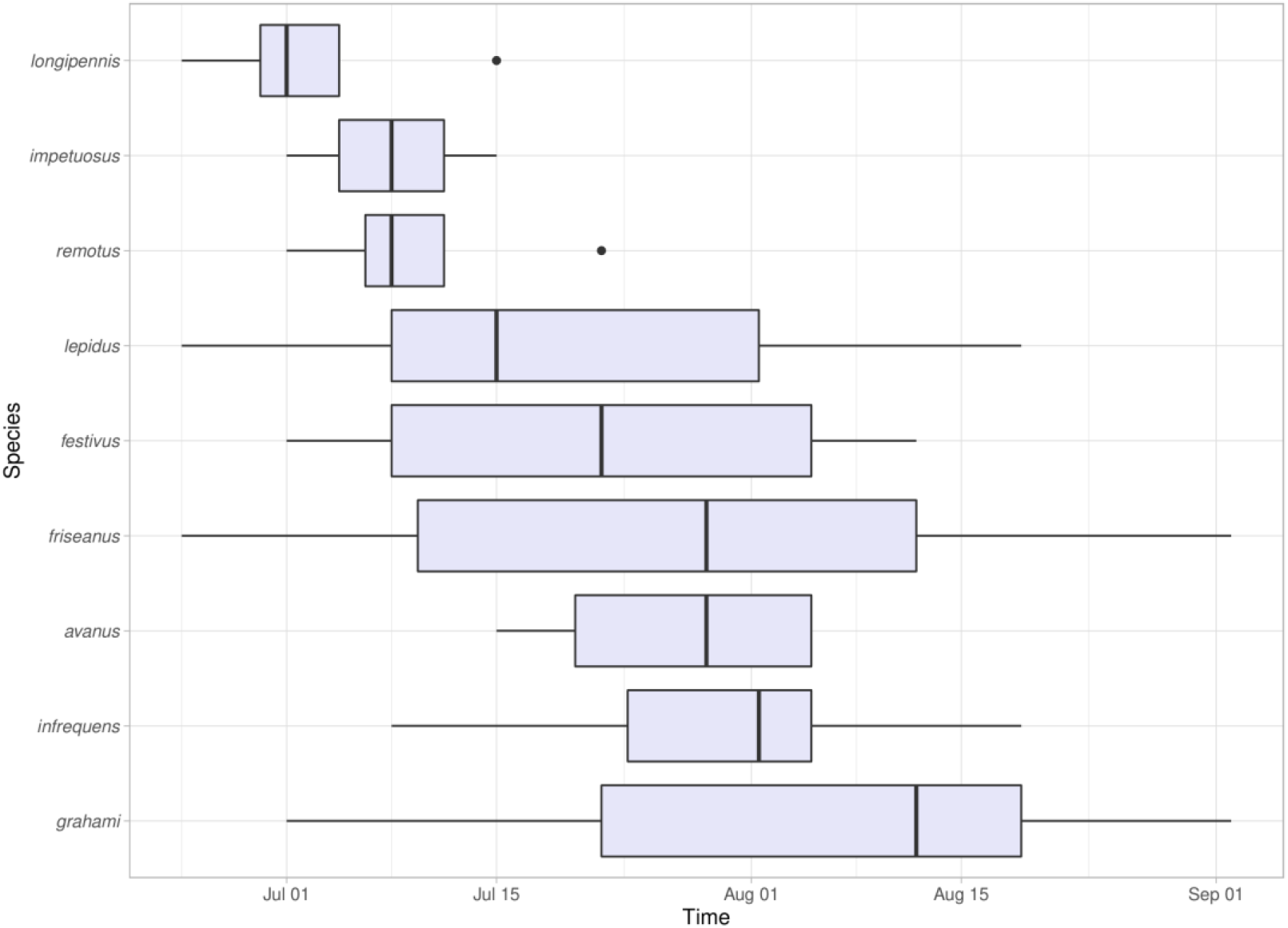
Relative periods of foraging activity of workers of nine *Bombus* species over summer months

### Spatial (Elevational) Patterns

Standardized abundances (Fig. 4) showed that the two most commonly collected species, *B. friseanus* and *B. lepidus,* were most common at middle elevations. Both species were less abundant at almost every other elevation. This can be contrasted with other less common species including *B. longipennis*, *B. avanus*, *B. festivus*, and *B. impetuosus* with narrower elevational distributions (Fig. 6).

## Discussion

There appear to be at least two factors maintaining bumblebee diversity on Yulong Snow Mountain. The first is temporal and based on the phenology of worker flights and foraging periods over the summer flowering season. The temporal difference in peak abundances of *B. friseanus* and *B. lepidus* suggests a partitioning effect separating the periods when each species is active (Fig. 3). Indeed, the weighted mean week of occurrence was more than three weeks earlier for *B. lepidus* than for *B. friseanus* (Table 2). We also see some temporal partitioning of less abundant species (Fig. 4). Previous studies on bumblebee diversity in other regions also show temporal partitioning. Lye et al. (2010) found evidence for temporal niche partitioning of bumblebees throughout the course of a day for four naturalized species in New Zealand. Pleasants (1980) found evidence of temporal niche partitioning of a bumblebee-dominated pollinator assemblage throughout the summer season in the Rockies of North America. Heinrich (1976) found clear partitioning of blooming periods for bumblebee-pollinated bog floras in Maine, USA, supporting the broader idea that many plant-pollinator assemblages minimize direct competition.

Spatial niche partitioning along an elevation gradient on Yulong Mountain appears to show its strongest influence among the rarer bumblebee species as they showed more limited distributions based on elevation. It is possible that, by remaining at lower elevations, these species avoid zones dominated by the two most common species. In contrast, *B. friseanus* and *B. lepidus* avoid some competition with each other through temporal partitioning as there appears to be little spatial partitioning along what appears to be a much broader elevational gradient. While *B. friseanus* was highly abundant at both 3200 m and 3400 m field sites, *B. lepidus* showed high abundance at 3400 m and lower abundances at other elevations including 3200 m (Fig. 5). There are clearly different specific site conditions for 3200 m and 3400 m leading to such differing abundances. Patterns of combined bumblebee abundance show a mid-elevational peak which parallels their thermoregulatory adaptations for lower ambient temperatures. Similar patterns have been shown for mammalian species on elevational gradients as well (McCain 2005).

**Fig. 5.**
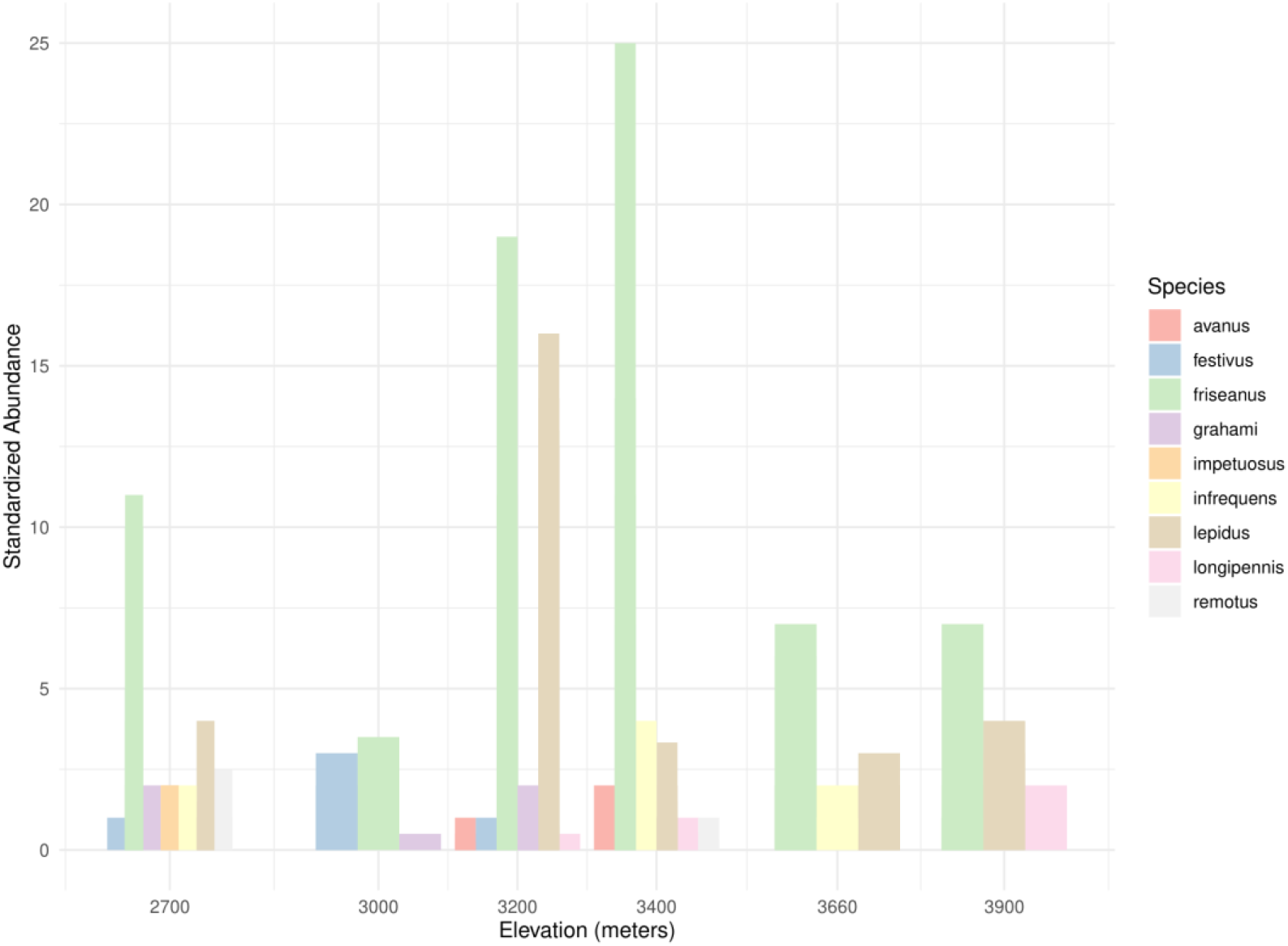
Distributions of nine *Bombus* species along an elevational gradient on the Yulong Snow Mountain

**Fig. 6.**
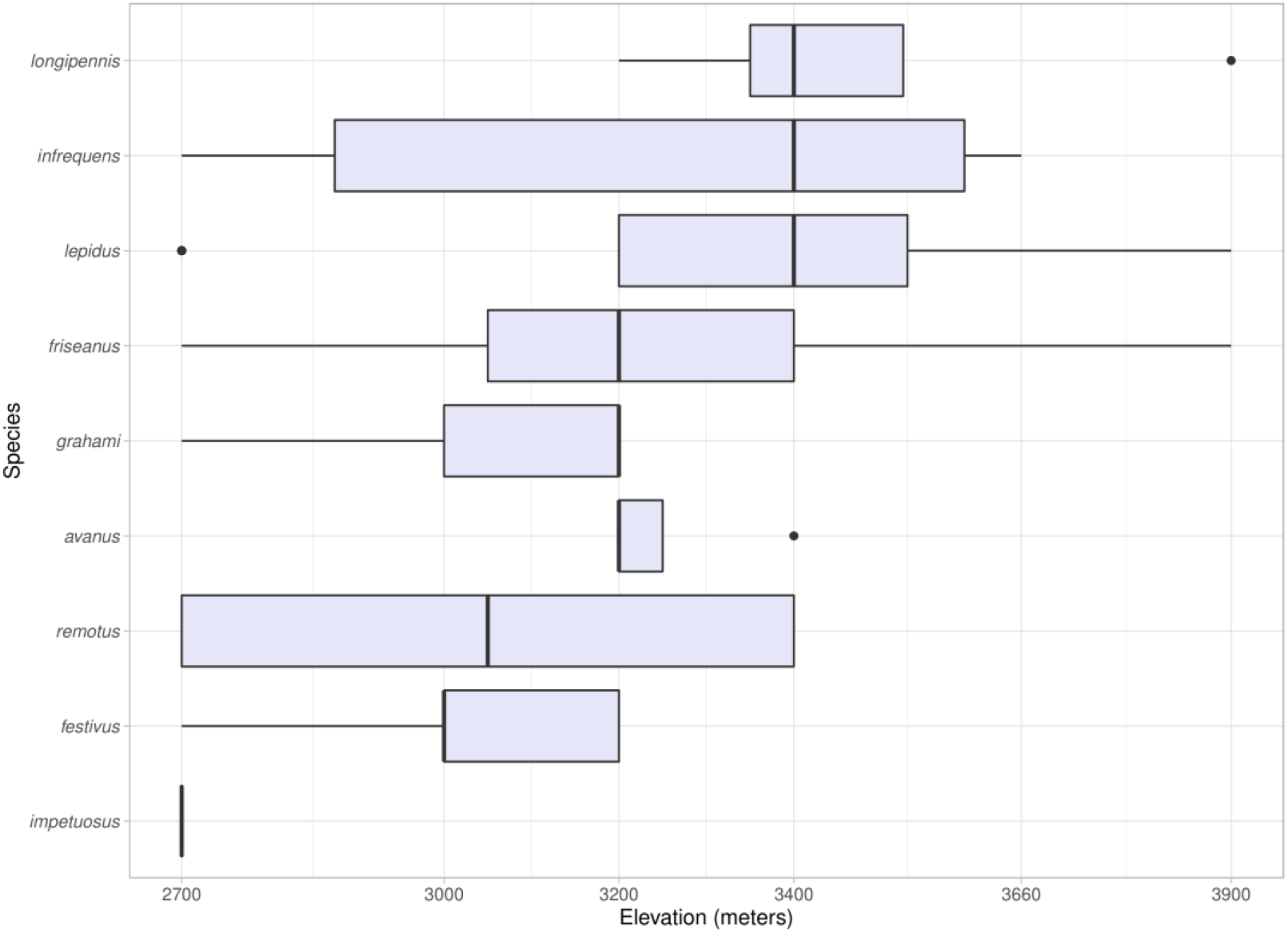
Worker abundance of nine *Bombus* species worker standardized for collection effort at different elevations along the elevational gradient of the Yulong Snow Mountain

In fact, our results are similar to findings in a report on bumblebees in the Himalayan range through Nepal (Williams et al. 2010) as latitude through the Nepalese region is similar to Yulong. The large elevational gradient and lengthy summer period of Yulong appears to contribute to bumblebee diversity on the mountain chain but may also be indicative of driving factors for bumblebee diversity throughout temperate elevations within the Himalayan chain. As elevational gradients produce many different niches for bumblebees this promotes specialization and the most common species are able to partition when they are most active preventing excessive competition.

This study covered 11-weeks from summer to early autumn but some bumblebee gynes begin emerging in Spring (April-May) in the Himalayas while *Rhododendron* spp. and other vernal angiosperms are in bloom. Future studies should sample the emergence of the earliest, vernal workers to the last, autumnal ones prior to winter. This would better show their interspecific phenological divergence and relative abundances would likely capture a greater bumblebee diversity. As *B. friseanus* showed peak relative abundance on week 11 of this study continuing to sample past week 11 would, in the future, offer a clearer picture of the phenological distribution of this common species.

Phenological mismatches caused by warming weather could also lead to the asynchronous emergence of bumblebee queens emerging from dormancy asynchronous with the bloom cycles of their primary food plants leading to population decline (Pyke et al. 2016). Spatial partitionings of bumblebee species could also be impacted by warming weather. Distributions of several species have been shown to change due to global warming as these organisms move to higher latitudes to maintain the temperate ranges they tolerate (Parmesan 2006). The same trend has been found in mountainous regions but the species are recorded moving to higher elevations instead (Marshall et al. 2020). Movement to higher elevations can lead to an increase in the potential for interspecific competition as more species overlap in smaller, geographic spaces. Species native to the highest elevations may now suffer from increasingly constricted distributions (Freeman et al. 2018). These bumblebees must either adapt to warmer temperature ranges or face extinction as they are “squeezed off” mountain tops. Furthermore, competition is compounded with phenological mismatches and/or floral resource mismatches due to differentiating rates of dispersal to higher elevations by winged bumblebees and plant propagules. Add invasive species, parasites/pathogens, pesticides, and land-use changes (Goulson et al. 2015) to global warming and you have the potential for creating a death-spiral for bumblebee diversity in Yunnan. This study then has the potential to be a snapshot of the current conditions on Yulong Snow Mountain that can be used for comparison in future studies as the mountain’s climate varies.

## Conclusions

Data analyses answers our original research questions. It affirms that temporal and spatial niche partitioning were two factors that helped maintain the diversity of worker bumblebees on Yulong Snow Mountain. Less abundant species were largely confined to lower elevations. The two, dominant species lowered competition over time as the abundance of workers of one species peaked earlier in the season compared to the second. This has important ramifications for the future because climate change has the potential to negatively impact on how temporal and spatial niche partitionings are sustained by affecting emergence times and species distributions.

## Acknowledgements

We are most grateful to Anne Beeman for her extensive help during field collections and specimen processing. Dr. Huan Liang, Mr. Xin Xu, Ms. Hai-Ping Zhang and Prof. Paul Williams are also thanked for their help identifying *Bombus* specimens. We thank Lijiang Forest Ecosystem Research Station on Yulong Snow Mountain for permission to conduct experiments and provide logistic support in the field.

