## Supplementary material for "Temporal and Spatial Niche Partitioning of Bumblebee Species (Apidae: *Bombus*) on a Himalayan Mountain": R Code and Datasets: Manuscript Code 1.R

```
# R Code for Temporal and Spatial Niche Partitioning Manuscript
# Created by Alan Moss

#package needed
library(ggplot2)
library(psych)

#import "Moss_1", "Moss_2", and "Moss_3" datasets first


### Figure 2
## Grouped barplot with smoothed mean line for all species over time
all.time.plot<-ggplot(Moss_2, aes(fill=Species,x=Week,y=Abundance.Sampling)) +
  geom_bar(position = "dodge", stat = "identity") +
  geom_smooth(se=FALSE,aes(color=Species,x=Week,y=Abundance.Sampling,group=Species)) + 
  scale_x_discrete(labels=c("w01" = "01", "w02" = "02", "w03" = "03","w04" = "04", "w05" = "05", "w06" = "06","w07" = "07", "w08" = "08", "w09" = "09","w10" = "10", "w11" = "11")) +
  scale_fill_brewer(palette="Pastel1") + 
  scale_color_brewer(palette="Set1") + ylab("Standardized Abundance") +
  theme(plot.margin = unit(c(2.5,2.5,2.5,2.5), "cm"))
all.time.plot + theme_light() + theme(plot.margin = unit(c(2.5,2.5,2.5,2.5), "cm"))


### Figure 3
## Comparison of B.friseanus and B.lepidus phenology 
# grouped barplot over 11 weeks
bar1<-ggplot(Moss_1, aes(fill=Species,x=Week,y=Abundance)) +
  geom_bar(position = "dodge", stat = "identity") +
  geom_smooth(se=FALSE,aes(color=Species,x=Week,y=Abundance,group=Species)) + 
  scale_fill_brewer(palette="Pastel1") + 
  scale_color_brewer(palette="Set1") +
  ylab("Standarded Abundance") +
  scale_x_discrete(labels=c("w01" = "01", "w02" = "02", "w03" = "03","w04" = "04", "w05" = "05", "w06" = "06","w07" = "07", "w08" = "08", "w09" = "09","w10" = "10", "w11" = "11")) +
  theme(plot.margin = unit(c(2.5,2.5,2.5,2.5), "cm"))
bar1 + theme_light() + theme(plot.margin = unit(c(2.5,2.5,2.5,2.5), "cm"))


### Figure 4
## boxplot based on time for all species
Moss_2$ï..Date <- as.Date(Moss_2$ï..Date,format = "%m/%d/%y")
Moss_2$Species <- factor(Moss_2$Species, levels=c("grahami","infrequens","avanus","friseanus","festivus","lepidus","remotus","impetuosus","longipennis"))
time.box<-ggplot(data=Moss_2, aes(x=Abundance.Sampling, y=ï..Date)) + geom_boxplot(aes(Species),  fill="lavender") + xlab("Species") + ylab("Time")
time.box + coord_flip()  + theme_light() +
  theme(axis.text.y = element_text(face = "italic"))  + theme(plot.margin = unit(c(2.5,2.5,2.5,2.5), "cm"))


### Figure 5
## grouped bar plot for all species over different elevations
elv.barplot<- ggplot(Moss_2, aes(fill=Species,x=Elevation,y=Abundance.Sampling)) +
  geom_bar(position = "dodge", stat = "identity") +
  scale_fill_brewer(palette="Pastel1") + 
  theme(plot.margin = unit(c(2.5,2.5,2.5,2.5), "cm")) +
  ylab("Standardized Abundance") + xlab("Elevation (meters)")
elv.barplot + theme_minimal() + theme(plot.margin = unit(c(2.5,2.5,2.5,2.5), "cm")) +
  scale_x_continuous(breaks=c(2700,3000,3200,3400,3660,3900))


### Figure 6
# boxplot for elevation for all species
Moss_2$Species <- factor(Moss_2$Species, levels=c("impetuosus","festivus","remotus","avanus","grahami","friseanus","lepidus","infrequens","longipennis"))
elv.boxplot <- ggplot(data=Moss_2, aes(x=Abundance.Sampling, y=Elevation)) + geom_boxplot(aes(Species), fill="lavender") + xlab("Species") + ylab("Elevation (meters)")
elv.boxplot + coord_flip() + theme(plot.margin = unit(c(2.5,2.5,2.5,2.5), "cm")) + 
  theme_light() + ylab("Elevation (meters)") + scale_y_continuous(breaks=c(2700,3000,3200,3400,3660,3900)) +
  theme(axis.text.y = element_text(face = "italic"))+ theme(plot.margin = unit(c(2.5,2.5,2.5,2.5), "cm"))
```
